# Activation of dorsal raphe serotonergic neurons promotes persistent waiting independently of reinforcing effects

**DOI:** 10.1101/012062

**Authors:** Madalena S. Fonseca, Masayoshi Murakami, Zachary F. Mainen

**Author notes:** To whom correspondence should be addressed, Address: Champalimaud Neuroscience Programme, Champalimaud Centre for the Unknown, Avenida de Brasilia, s/n, 1400-038, Lisbon, PORTUGAL.

## Abstract

The central neuromodulator serotonin (5-HT) has been implicated in a wide range of behaviors and affective disorders, but the principles underlying its function remain elusive. One influential line of research has implicated 5-HT in response inhibition and impulse control. Another has suggested a role in affective processing. However, whether and how these effects relate to each other is still unclear. Here, we report that optogenetic activation of 5-HT neurons in the dorsal raphe nucleus (DRN) produces a dose-dependent increase in mice’s ability to withhold premature responding in a task that requires them to wait several seconds for a randomly delayed tone. The 5-HT effect had a rapid onset and was maintained throughout the stimulation period. In addition, movement speed was slowed but stimulation did not affect reaction time or time spent at the reward port. Using similar stimulation protocols in place preference and value-based choice tests, we found no evidence of either appetitive or aversive effects of DRN 5-HT neuron activation. These results provide strong evidence that the efficacy of DRN 5-HT neurons in promoting waiting for delayed rewards is independent of appetitive or aversive effects and support the importance of 5-HT in behavioral persistence and impulse control.

## INTRODUCTION

The central neuromodulator serotonin (5-HT) has been implicated in a variety of different sensorimotor, affective and cognitive behaviors. However, its precise role in specific behaviors and the general principles that underlie its diverse effects, remain unclear. 5-HT function has long been associated with response inhibition and impulse control [1, 2] based largely on evidence that reductions in 5-HT levels, produced by lesion of 5-HT neurons in the dorsal raphe nucleus (DRN), the chief source of serotonin to the forebrain, tryptophan depletion, or pharmacological blockade, dis-inhibit behavior suppressed by punishment [3–10] (for review, see [1]) and increase premature responding for rewards [11–15] (for review, see [16]). In a recent series of studies, Miyazaki and colleagues showed that neurons in the DRN were active while rats waited for delayed rewards and delayed reward-predictive cues [17], that local pharmacological blockade of DRN serotonergic activity increased premature responding [18], and that optogenetic activation of DRN 5-HT neurons decreased it [19]. These findings led to the more specific proposal that 5-HT facilitates waiting to obtain rewards, promoting “patience” [20].

Serotonin has also been implicated in behaviors involving both appetitive (rewarding) and aversive (punishing) stimuli. On the one hand, serotonin has been linked to positive affect pharmacologically, via the anti-depressant effects of selective serotonin reuptake inhibitors (SSRIs) and the ‘empathogenic’ effects of 3,4-(±)-methylenedioxymethamphetamine (MDMA) [21]. DRN neurons have been shown to respond to reward-related stimuli [22–25] and reductions in 5-HT by tryptophan depletion have been shown to affect reward processing [26–28]. On the other hand, 5-HT has also been linked to aversive processing [29–32]. Serotonergic neurons are activated by aversive stimuli such as electric shocks [33, 34] and pharmacological increases in 5-HT attenuate the effects of medial frontal bundle stimulation [35], dopamine-dependent response potentiation [36] and painful mechanosensory stimuli [37, 38]. Attempting to synthesize some of these diverse results, a prominent computational theory has proposed 5-HT as an opponent process to dopamine, in mediating prediction errors [39, 40].

These lines of evidence raise the question of whether and how the 5-HT involvement in response inhibition might be related to its appetitive and aversive effects. For example, 5-HT could be involved specifically in the withholding of responses that would otherwise lead to punishment [10, 32, 40–42]. Analogously, a positive affective function of 5-HT could explain enhanced waiting, by, for example, inducing a state that animals seek to maintain. In support of the latter possibility, Liu *et al*. [43] showed that optogenetic stimulation of DRN 5-HT neurons can serve as an appetitive reinforcer in several behavioral tasks, including spatial preference tests and explicit choice tasks. Miyazaki *et al.* [19] saw no such appetitive effects of DRN 5-HT neuron stimulation, but using a different optogenetic protocol and reward assay. Thus, the differences between the two studies leave unclear whether or not 5-HT effects on waiting are accompanied by, or even a consequence of, some form of appetitive signal. One possibility is that the more transient and synchronous stimulation employed by Liu *et al.* [43] produces different effects from the more sustained stimulation employed by Miyazaki *et al.* [19], much as phasic and tonic dopamine have been shown to have different effects [44, 45].

To address these issues, in the present report, we performed a series of experiments examining the effects of optogenetic activation of DRN 5-HT neurons in both a waiting task and in a series of assays that assess value via preference and choice. We found that photostimulation of DRN 5-HT neurons led to a dose-dependent prolongation in the ability of mice to withhold premature responding in a waiting task. In contrast, using the same photostimulation parameters, we found no evidence of an appetitive or aversive effect of DRN 5-HT activation in two place preference tests, and a third, extremely sensitive, value-based choice task. These results provide direct evidence of sufficiency of DRN 5-HT activation in promoting waiting behavior, providing definitive evidence for a link between 5-HT and behavioral changes that is independent of reinforcing effects.

## RESULTS

### Waiting behavior

To study the effect of DRN 5-HT stimulation on waiting behavior, we trained adult transgenic mice expressing CRE recombinase under the serotonin transporter promoter (SERT-Cre) and wild-type littermates (WT) in a waiting task (**Fig. 1** and Experimental Procedures) until their performance was stable (minimum 2 months, see Experimental Procedures). The experimenters were blind to mice’s genotype throughout training, surgery and testing. The task required mice to wait for a randomly delayed tone (exponential distribution, min 0.5s, mean: 1.5 – 16 s) (**Fig. 1A** inset) in order to obtain a water reward at the reward port. The mean value for the tone delay was adjusted for each animal so that it successfully waited in 50 – 60% of the trials. This allowed us to detect both possible increases and decreases in waiting time caused by DRN 5-HT stimulation. The random delay promoted a wide distribution of waiting times (**Fig. 1B**, **C**) [46] providing a sensitive measure of waiting and enabling us to study the dynamics of stimulation effects. Mice performed around 423.2 ± 122.3 trials per session, mean ± S.D., total N = 128 sessions from 11 mice).

**Figure 1.**
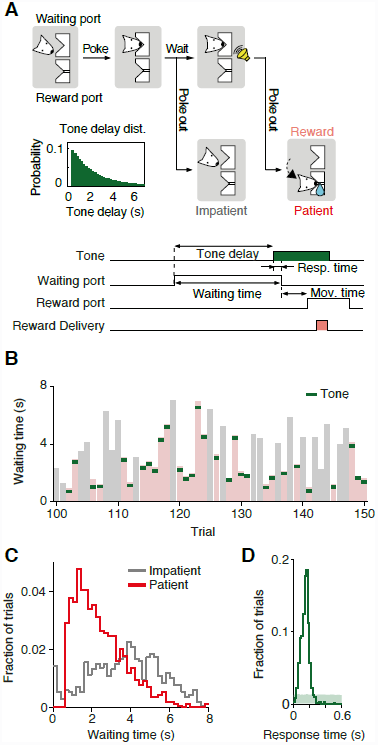
The waiting task and the behavioral performance. **(A)** Schematic diagram of trial events in the waiting task (top). In each trial, a mouse is required to wait for a randomly delayed tone and move to the reward port to obtain a water reward (Patient trial). If the mouse fails to wait for the tone, the reward is not available (Impatient trial). An example of the probability distribution of the delays to the tone is shown in the inset. Timeline of the task events and definition of the behavioral parameters (bottom). The green rectangle indicates presentation of the tone and the pink rectangle indicates the water reward. **(B)** Snapshot of the waiting behavior. The waiting period in each trial is indicated as a light red or gray bar, representing patient and impatient trials, respectively. Green ticks represent the presentation of the tone. **(C)** Waiting time histograms of impatient trials (gray) and patient trials (red) of an example mouse. The histograms show data pooled across sessions. **(D)** A histogram of response time to the tone of an example mouse. The light shaded area indicates 95% range of response time histograms from the shuffled data.

We classified the trials into two types: those in which the mouse waited for the tone (‘patient trials’, 53.7 ± 6.3%, mean ± S.D., N = 11 mice) and those in which the mouse exited the waiting port before the presentation of the tone (‘impatient trials’). The distributions of waiting time for patient and impatient trials from a representative mouse are shown in **Figure 1C**. The median waiting time ranged from 1.0 – 7.5 s across mice, with a mean of 2.8 ± 1.8 s (mean ± S.D.). Mice understood the tone-reward association, as shown by the prompt response to the tone (**Fig. 1D**, median response time 150 ms for the example mouse, population range: 74 – 316 ms).

### Optical activation of DRN 5-HT neurons during waiting

We used optogenetic methods to selectively activate DRN 5-HT neurons in the waiting task. After training, we expressed the light-sensitive ion channel channelrhodopsin-2 (ChR2) in DRN 5-HT neurons using an AAV2/1 viral vector (AAV-Dio-ChR2-EYFP) injected into the DRN of SERT-Cre mice (N = 7) or wild-type littermates (WT, N = 4). An optical fiber was then implanted in the same location. Both SERT-Cre and WT animals were infected, implanted and stimulated in the same manner (**Fig. 2A**) (see [38] for more details about the targeting strategy and validation). Histology performed at the end of testing showed ChR2-YFP expression localized to the DRN in SERT-Cre animals (**Fig. 2B**) and no expression in WT controls (not shown).

**Figure 2.**
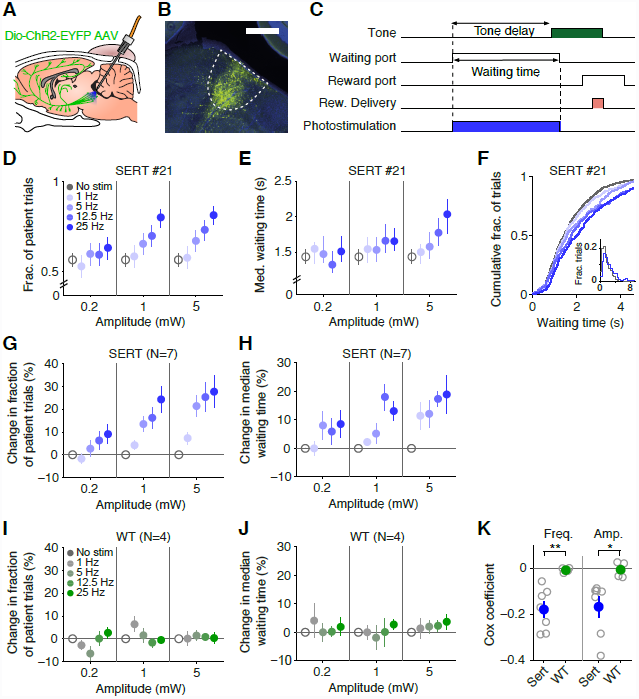
Photostimulation of DRN 5-HT neurons promotes waiting behavior. **(A)** A schematic drawing of the optogenetic approach. DRN neurons are infected with AAV1/2-Dio-ChR2-YFP. In SERT-Cre mice, 5-HT neurons will express ChR2-YFP (green cells) and can be photoactivated with blue light delivered through an optical fiber implant. **(B)** Fluorescence picture of a parasagittal section showing localized ChR2-YFP expression in the DRN. YFP in green. DAPI in blue. Scale bar, 500 m. **(C)** Photostimulation period (blue rectangle) is shown along with the task events. The same format as in **Figure 1A**. **(D)** Dose-dependent increase in the fraction of patient trials with DRN 5-HT stimulation from an example mouse, SERT #21. Error bars indicate 95% confidence intervals estimated with binomial fitting. Note that all the non-stimulated trials are pooled together and repeatedly plotted across the three amplitudes for visualization purpose, here and elsewhere. **(E)** Dose-dependent increase in median waiting times with DRN 5-HT stimulation in the same example mouse. Error bars indicate a 95 percentile range (2.5 – 97.5 percentile) of a bootstrap distribution. **(F)** Cumulative histograms of waiting times across frequencies (at 5mW) from the same mouse. Inset shows waiting time histograms of non-stimulated and stimulated (25 Hz at 5mW) trials. Patient and impatient trials are pooled together in each histogram. **(G)** Dose-dependent increase in the fraction of patient trials for the population of SERT-Cre mice (N = 7). Percent changes in the fraction of patient trials with respect to non-stimulated trials, averaged across mice, are shown. Error bars indicate S.E.M.. **(H)** Dose-dependent increase in median waiting times for the population of SERT-Cre mice (N = 7). Percent changes in median waiting time with respect to non-stimulated trials, averaged across mice, are shown. Error bars indicate S.E.M.. **(I)** The same as **G** but for the population of wild-type mice (N = 4). **(J)** The same as **H** but for a population of wild-type mice (N = 4). **(K)** Cox regression, demonstrating a dose-dependent increase in waiting performance. Cox regression coefficients for both frequency (left) and amplitude (right) are shown for SERT-Cre mice (blue) and wild-type mice (green). Individual mice in open circles. Averages across mice are shown in filled circles. Error bars represent S.E.M.. **: *P* < 0.01; *: *P* < 0.05 with independent two-sample t-test (SERT v.s. WT). Note that negative coefficients indicate that the higher the frequency (or amplitude), the lower the leaving rate, thus the longer the waiting time.

To test the effect of DRN 5-HT activation on waiting behavior, we delivered photostimulation during the waiting period (from waiting port entry to waiting port exit, see **Fig. 2C**). In order to obtain a full dose-response curve, mice were stimulated with different frequencies (0, 1, 5, 12.5, 25 Hz) and amplitudes (0.2, 1, 5 mW) in randomly interleaved trials. Testing began 3 – 4 weeks after the surgery to allow for virus expression and re-training. This experiment consisted of data from 10 – 16 sessions per mouse (excepting one mouse that contributed only 4 sessions before losing the implant). No data was excluded.

Photostimulation in SERT-Cre mice, but not WT littermates, resulted in an increase in the fraction of patient trials and median waiting time. **Figure 2** shows waiting task performance for a representative SERT-Cre mouse **(Fig. 2D-F)** and the population of SERT-Cre mice (**Fig. 2G**,**H**) and WT mice (**Fig. 2I**,**J**). These effects were confirmed with a 3-way ANOVA (frequency, amplitude, genotype) on the normalized fraction of patient trials (main effect of genotype, F_(1,9)_ = 10.220, *P* = 0.011, frequency X genotype, F_(2.205, 19.845)_ = 3.392, *P* = 0.002, amplitude X genotype: F_(1.566, 14.097)_ = 3.913, *P* = 0.053, no other terms involving genotype were significant) and normalized median waiting time (frequency X genotype: F_(3.000,27.000)_ = 4.926, *P* = 0.007, amplitude X genotype: F_(2.000,18.000)_ = 3.565, *P* = 0.050, no other terms involving genotype were significant), followed by separate 2-way ANOVAs restricted to SERT (fraction of patient trials: main effect of frequency, F_(1.718,10.307)_ = 16.439, *P* = 0.001, main effect of amplitude: F_(1.353,8.118)_ = 10.633, *P* = 0.008, N = 7 mice; median waiting time: main effect of frequency, F_(3,18)_ = 10.094, *P* < 0.001, main effect of amplitude: F_(2,12)_ = 8.303, *P* = 0.005, N = 7 mice) and WT (no significant effects in either measure, N = 4 mice).

The dose-dependency of stimulation frequency and amplitude on waiting time, were confirmed by a Cox regression. Cox regression is a form of survival analysis suited for time-to-event data and that allows one to quantify the effect of several predictor variables (i.e. photostimulation frequency and amplitude) on the time it takes for a specific event (i.e. giving up waiting) to occur (see Experimental Procedures for more details on the model). The analysis yielded a negative coefficient for both frequency and amplitude in SERT-Cre animals (**Fig. 2K**; frequency: –0.179 ± 0.035; amplitude: –0.167 ± 0.044, mean ± S.E.M., N = 7 mice), that was significantly different from zero (one-sample t-test for frequency: *P* = 0.002, N = 7 mice, and amplitude: *P* = 0.009, N = 7 mice) and significantly different from WT (two-sample independent t-test for frequency (SERT, WT), *P* = 0.005; and amplitude, *P* = 0.0267, N_SERT_ = 7, N_WT_ = 4), demonstrating a frequency- and amplitude-dependent reduction in the probability of giving up waiting and thus, an increase in waiting time (see Experimental Procedures for details).

### Time course of the waiting effect

To characterize the time-course of the photostimulation effect on waiting, we estimated the hazard rate of leaving the waiting port across time. The hazard rate measures the rate at which the mouse left the waiting port given that it had successfully waited to that point, as a function of time spent waiting (see Experimental Procedures for details). Hazard rate for a representative SERT-Cre mouse is shown in **Figure 3A** and the average across the SERT-Cre population in **Figure 3C**. DRN 5-HT photostimulation led to a reduction in the hazard rate of leaving in a manner that was both frequency-dependent (**Fig. 3A**, see **Fig. 2K** for Cox regression coefficients) and amplitude-dependent (not shown, see **Fig. 2K** for Cox regression coefficients). Inspection of the hazard rate plots (**Fig. 3A**, **C**) suggests that hazard rate is reduced in the first time bins and that the difference lasts throughout the waiting period (i.e. the effect does not decrease or reverse over stimulation time). To estimate how early the effect of photostimulation could be detected we compared the hazard rates for trials with and without photostimulation. **Figure 3B** shows the cumulative hazard for an example mouse. To estimate the earliest detectable effect we calculated the time at which the two cumulative curves could be distinguished by a permutation test (see Experimental Procedures for details). Five of the seven SERT-Cre mice had detectable onset times below 1 s (**Fig. 3D**, range: 0.50 – 2.14 s; population median 0.66 s).

**Figure 3.**
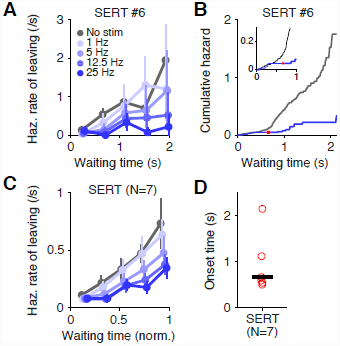
Time course of the effect of DRN 5-HT photostimulation. **(A)** Fast onset and sustained effect of DRN 5-HT stimulation. Hazard rate of leaving the waiting port is plotted across time for an example mouse, SERT #6. Only data from the 5mW photostimulation conditions is shown. **(B)** Fast onset of the effect of DRN 5-HT photostimulation. Cumulative hazard rate is plotted across time. Data with non-stimulation and 25 Hz at 5mW stimulation trials are shown. The red circle indicates the detectable onset time. The inset shows the zoomed-in view near the origin. **(C)** Hazard rate of leaving for the population of SERT-Cre mice (N = 7). Averages across mice are shown. Error bars indicate S.E.M.. **(D)** Detectable onset times across the population of SERT-Cre mice (N = 7). The black bar indicates the average across mice. Individual mice are shown in open circles.

### Effect of DRN 5-HT neuron optical stimulation on other task periods

One possibility consistent with the increase in waiting task performance is that photostimulation has a general motoric effect. To investigate this possibility, we first analyzed the movement time (time taken from exiting the waiting port to entering the reward port) in the same task and sessions as above (**Fig. 4A**). Median movement time was increased following photostimulation of DRN 5-HT neurons, shown for an example mouse (**Fig. 4B**, only 5mW conditions are shown; No-stim., 424ms; Stim. (25Hz at 5mW), 450ms; 6.03% change) and population (**Fig. 4C**, 9.55 ± 1.84% change for Stim. (25Hz at 5mW), mean ± S.E.M., N = 7 mice). The effects on movement time were significant at the population level in a 3-way ANOVA including genotype as a factor (Frequency x genotype: F_(3.000,27.000)_ = 11.280, *P* < 0.001, Amplitude x genotype: F_(1.778,15.999)_ = 8.631, *P* = 0.004) and a follow-up 2-way ANOVA restricted to SERT-Cre (Frequency, F(2.652,15.911) = 16.851, *P* < 0.001, Amplitude, F(1.432,8.594) = 17.117, *P* = 0.002, Frequency x Amplitude, F(4.194,25.165) = 6.177, *P* = 0.001, N = 7 mice) and WT (no significant effects, N = 4 mice).

**Figure 4.**
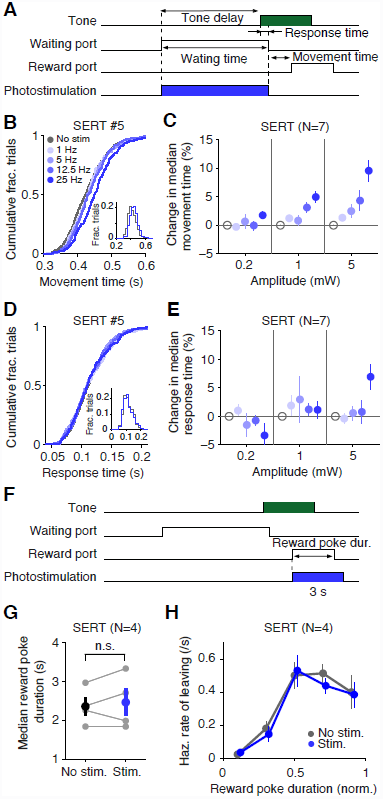
The effect of DRN 5-HT photostimulation on movement time, response time and reward poke duration. **(A)** Photostimulation period (blue rectangle) and the definition of behavior parameters are shown along with the task events. The same format as in **Figure 1A**. **(B)** Cumulative histograms of movement times across frequencies (at 5 mW) from an example mouse, SERT #5. Inset shows movement time histograms of non-stimulated and stimulated (5mW, 25 Hz) trials. **(C)** Increase in movement times with DRN 5-HT stimulation for the population of SERT-Cre mice (N = 7). Percent changes in movement times with respect to non-stimulated trials, averaged across mice, are shown. Error bars indicate S.E.M.. **(D)** Cumulative histograms of response times across frequencies (at 5 mW) from an example mouse, SERT #5. Inset shows response time histograms of non-stimulated and stimulated (25Hz at 5mW) trials. **(E)** No systematic changes in response times with DRN 5-HT stimulation for the population of SERT-Cre mice (N = 7). Percent changes in response times with respect to non-stimulated trials, averaged across mice, are shown. Error bars indicate S.E.M.. **(F)** Reward port stimulation experiment. A subset of the animals used in the waiting period stimulation experiment (SERT-Cre, N = 4; WT, N = 2, not shown). Photostimulation period (blue rectangle) and the definition of the behavior parameter are shown along with the task events. The same format is used as in **Figure 1A**. **(G)** No significant change in reward poke duration with DRN 5-HT stimulation for the population of SERT-Cre mice (N = 4). Individual mice are shown in gray circles. Averages across mice are shown in filled circles. Error bars indicate S.E.M.. **(H)** No significant change in hazard rate of leaving the reward port with DRN 5-HT stimulation, averaged across mice. Error bars indicate S.E.M..

In contrast, we found that the response time (the time between tone onset and movement onset, **Fig. 4A**) was not affected in the same mouse (**Fig. 4D**, only 5mW conditions are shown; No-stim., 112ms; Stim. (25Hz at 5mW), 111 ms; –1.16% change) and there were no significant effects on response time at the population level (**Fig. 4E**, 6.92 ± 0.02% change for Stim. (25Hz at 5mW), N = 7 mice), as indicated by a 3-way ANOVA (minimum *P* = 0.175). Although the strongest stimulation parameter (5 mW, 25 Hz) produced apparent increase in the response time (Wilcoxon signed-rank test, *P* = 0.023), it did not survive correction for multiple comparisons (Bonferroni correction). Note that the response time period is included in the waiting time measurement and occurs when photostimulation is active, whereas the movement time period occurs after the photostimulation has turned off.

To investigate this issue further, we ran a separate experiment in which stimulation was delivered while animals were at the reward port (25 Hz at 5 mW, 0 – 3 s from port entry) (**Fig. 4F**). Similar to the waiting period, the reward period also required the mice to ‘poke-in’ at a nose port and was thus very similar to waiting in terms of motor parameters. However, photostimulation had no significant effect on the median reward poke duration (**Fig. 4G**, paired t-test, *P* = 0.528, N = 4 mice) or on the hazard rate of leaving (**Fig. 4H**, Cox coeff. –0.1087 ± 0.1125, one-sample t-test, *P* = 0.405, N= 4 mice). These results confirm that there is behavioral specificity in the effects of DRN 5-HT stimulation.

### Conditioned and real-time place preference

To investigate whether the increase in waiting times observed was related to possible appetitive or aversive reinforcing effects of DRN 5-HT photostimulation [43], we tested the ability of photo-simulation to induce conditioned place preference (CPP), a standard assay for testing the rewarding effects of natural and artificial stimuli (**Fig. 5A**, day 1 – 4). After a habituation session (pre-test), mice were subjected to two days of conditioning (day 2 – 3). On each day, mice were confined to one side of the box and given either photostimulation (12.5 Hz at 5 mW, for 3 s, every 10 s) or no stimulation. They were then placed in the other side chamber and given the alternative treatment. On day 4, mice were re-tested in the absence of stimulation (post-test). Occupancy plots (time spent in each region of space) for pre- and post-test for an example mouse are shown in **Figure 5B**. Preference for the chamber paired with photostimulation was assessed using a preference score calculated as the difference between the times spent in the stimulated and non-stimulated sides, divided by the sum of the two. We found no difference in preference score between pre- and post-test (**Fig. 5C**, paired t-test, *P* = 0.609, N = 4 mice) nor between SERT-Cre and WT controls (**Fig. 5D**, two-sample independent t-test, *P* = 0.728, N = 4 per group).

**Figure 5.**
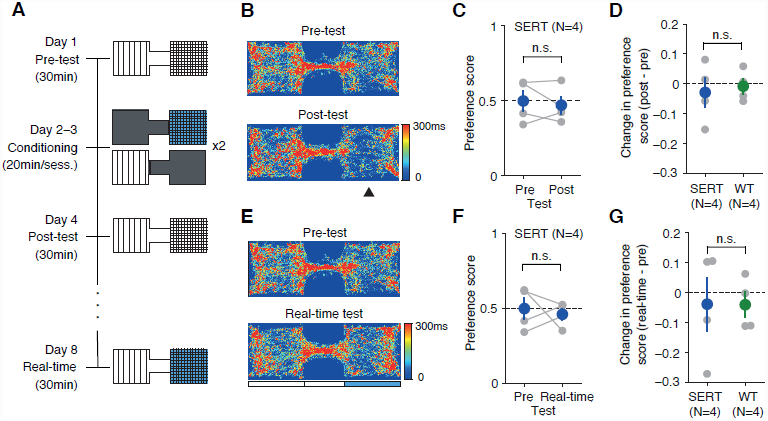
The effect of DRN 5-HT photostimulation in the place preference tests. **(A)** Conditioning and testing schedule for the conditioned place preference (CPP) test (Day 1 – 4) and real-time place preference test (Day 8). **(B)** Occupancy plots for the Pre-test session and Post-test session for an example SERT-Cre mouse, SERT #28. The photostimulation paired chamber is indicated by the arrowhead. **(C)** No significant change in the preference score after conditioning. Individual mice shown in gray circles. Averages across mice in blue. Error bars indicate S.E.M.. n.s.: not significant. **(D)** No significant difference in the change in the preference score between SERT-Cre and WT mice for the conditioned place preference test. Individual mice shown in gray circles. Average across mice shown in blue (SERT-Cre) or green (WT). Error bars indicate S.E.M.. **(E)** Occupancy plots for Pre-test session and real-time stimulation session for an example SERT-Cre mouse, SERT #28. The blue rectangle indicates the photostimulation-associated chamber. Note that the pre-test session is common for both the conditioned place preference test and real-time preference test. **(F)** No significant difference in the preference score between the pre-test and real-time session. The same format as shown in **C**. **(G)** No significant difference in the change in the preference score between SERT-Cre and WT mice for the real-time place preference test. The same format as shown in **D**.

We then tested the same mice on a real-time version of the place preference test, four days later. In this test, entry to the stimulation-assigned chamber (the same as the stimulation paired side in the CPP test) triggered optical stimulation (12.5 Hz at 5 mW for 3 s, every 10 s, until chamber exit). This version of the place preference procedure is more similar to the waiting time task in that mice could increase the time of stimulation by staying in one of the chambers. Occupancy plots for an example mouse are shown in **Fig 5E**. Consistent with the results in the standard CPP assay, there was no significant difference in preference between pre-test and real-time test (**Fig. 5F**; paired t-test, *P* = 0.701, N = 4 mice) or between SERT-Cre and WT animals (**Fig. 5G**; independent two-sample t-test, *P* = 0.987, N = 4 per group). Thus, DRN 5- HT stimulation showed no reinforcing effects, appetitive or aversive in either standard or real-time place preference tests.

### Probabilistic reward task

To further investigate the possible reinforcing effects of DRN 5-HT stimulation, we also trained and tested mice (the same used in the conditioned and real time place preference) in a probabilistic reward task that elicits reward probability matching behavior (**Fig. 6**) [47–49]. We chose this task for two main reasons. First, the probabilistic nature of the water reward schedule used in the task, by promoting probabilistic choice behavior, makes the choice susceptible to possible small biasing effects of photostimulation. Second, the availability of thousands of trials increases statistical power for detecting small possible stimulus effects.

**Figure 6.**
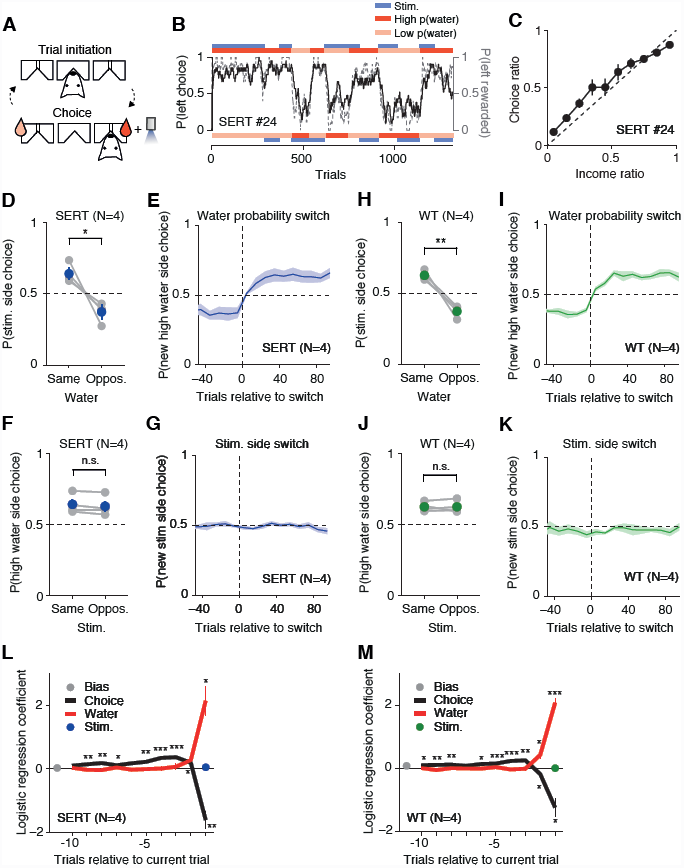
The effect of DRN 5-HT photostimulation in the probabilistic reward task. **(A)** Schematic diagram of trial events in the probabilistic reward task. In each trial, a mouse is required to enter the center port then move to the reward port to obtain a water reward delivered in a probabilistic manner. The pink and red water drops indicate the side port associated with the lower and higher probability of water reward, respectively. The blue light indicates the side port associated with the photostimulation. **(B)** The block schedule and mouse choice behavior from example sessions. Probability of choosing the left port (black solid line) overlaid with probability of obtaining reward at the left port (gray dashed line) (moving average of past 20 trials) are shown across trials for an example mouse, SERT #24. Two example sessions are concatenated. The top red/pink bar indicates the probability of water reward associated with the left port in a block of trials. The top blue bar indicates blocks where the left port was associated with the photostimulation. The bottom bars represent the same but for the right port. **(C)** Matching behavior. The choice ratio (probability of choosing left) is plotted as a function of income ratio (fraction of obtained reward at the left port in the past 20 trials) for the same mouse. The dashed line indicates the unity line. **(D)** Strong effect of water probability on choice. Probability of choosing the stimulation associated side is plotted separately for the blocks in which the higher water probability was assigned to the same side as the stimulation side and for the blocks in which the higher water probability was assigned to the opposite side. Individual SERT-Cre mice are shown in gray. Average across mice shown in blue. Error bars indicate S.E.M.. *: *P* < 0.05 with paired t-test **(E)** Prompt switch in the choice behavior in response to the change in the water probability. Probability of choosing the higher water probability side after the block switch (lower water probability side before the switch) is plotted across trials (bin = 10 trials) aligned on the block switch, for SERT-Cre mice (N = 4). Only the block switches in which the water probability changed without a change in stimulation side are included. The shaded area indicates S.E.M.. **(F)** No significant effect of DRN 5- HT stimulation on choice. Probability of choosing the higher water probability side is plotted separately for the blocks in which the photostimulation was assigned to the same side as higher water side and for the block when the photostimulation was assigned to the opposite side. The same format as in **D**. n.s.: not significant. **(G)** No apparent shift in choice behavior in response to the change in the stimulation side. Probability of choosing the stimulation side after the block switch (no stimulation side before the switch) is plotted across trials aligned on the block switch for SERT-Cre mice (N = 4). Only block switches in which the stimulation side changed without a change in the water probability are included. The shaded area indicates S.E.M.. **(H-K)** The same as **D-G** but for wild type littermates (WT, N = 4). **(L, M)** No significant effect of photostimulation on choice with a logistic regression analysis. **(L)** Logistic regression coefficients are plotted for the trial history variables up to the 10th trial back for SERT-Cre mice. Filled circles and thick lines indicate population mean. Error bars indicate S.E.M.. In some cases, the error bars are too small to be visible.***: *P* < 0.001, **: *P* < 0.01, *: *P* < 0.05. **(M)** The same as **L**, but for WT littermates.

The task required mice to initiate the trial with a nose poke entry and then choose one of two choice ports (**Fig. 6A**, **B**). Each choice port was associated with a specific reward probability (40 or 10%) and photostimulation probability (100 or 0%) for the length of a block (50 to 150 trials, randomly chosen). After completion of one block, new reward and stimulation probabilities were randomly chosen for the next block. Reward and stimulation probabilities were never the same in the two choice ports. If a reward was assigned to a choice port, and that port was not chosen, the water reward remained available until that side was chosen.

**Figure 6C** shows the fraction of choices made to one of the choice ports (choice ratio) as a function of the fraction of rewards the animal received from that port (income ratio) over all sessions of one example mouse. The observed behavior (**Fig. 6C**) generally conformed to the prediction from Herrnstein’s matching law [47], but showed a slight tendency towards under-matching (slope less than unity, also seen in the rest of the animals), as commonly observed [48]. In our experiment, under-matching would facilitate the detection of a superimposed biasing effect of photostimulation.

The impact of water rewards on behavior is clear from **Figure 6E**, where it can be seen that mice’s choice probability tracked switches in water probabilities (while stimulation probabilities remained constant). Overall, as expected, there was a significant effect of water rewards on overall choice probabilities: the probability of choosing the stimulation side in blocks where stimulation and high water probability were on the same side was significantly higher than when they were on opposite sides (water, paired t-test, *P* = 0.027, N = 4 mice).

In contrast, using the same analysis methods, we found no significant difference between the probability of choosing the “high water probability side” in blocks where photostimulation was present on that side or on the opposite side (**Fig. 6F**; paired t-test, *P* = 0.087, N = 4 mice). Similarly, there was no evidence that mice’s choice behavior tracked switches in stimulation side. Similar results were obtained for WT littermate controls injected, implanted and tested in the same manner (**Fig. 6H-K**; water: *P* = 0.005, stim: paired t-test, *P* = 0.982, N = 4 mice).

We further analyzed the effect of photostimulation on choice behavior using a logistic regression analysis [49] (see Experimental Procedures). Briefly, we generated a logistic regression model predicting the choice behavior using the past trial history, including choice history, water reward history and photostimulation history. We asked whether the past photostimulation history (and also other factors) had a significant effect on the choice in the current trial. As shown in **Figure 6L**, although the past water reward history and choice history had a strong effect on the choice behavior, we saw no effect of photostimulation history on choice (one-sample t-test, SERT vs. 0, *P* = 0.173, N = 4 mice). This was also true for WT animals **(Fig. 6M**, one-sample t-test, WT vs. 0, *P* = 0.815, N = 4 mice). These results are consistent with the results seen in conditioned and real-time place preference.

Finally, to exclude the possibility that the lack of reinforcing effects of photostimulation was due to insufficient ChR2 expression or some other difference relative to the mice tested in the waiting task, we modified the probabilistic reward task (**Fig. 7A**) so that the mice had to wait at the reward port for a variable delay to obtain a water reward (min 0 s, mean selected for each session to achieve around 50% waiting success; see Experimental Procedures for details). We expected photostimulation to increase patient waiting but based on the results of Miyazaki *et al.* [19] that these effects would be smaller than those during tone waiting. Photostimulation indeed led to a reduction in the hazard rate of leaving the reward port while waiting for a delayed reward (**Fig. 7B**, **C**). This was confirmed by a significant reduction in Cox coefficient compared to WT mice (**Fig. 7D**, two sample independent t-test, SERT vs. WT, *P* = 0.04), although this effect did not reach statistical significance in SERT-Cre mice alone (one sample t-test, SERT vs. 0, *P* = 0.08). Thus, despite having undergone a long period of expression and testing photostimulation was still effective in producing waiting time effects.

**Figure 7.**
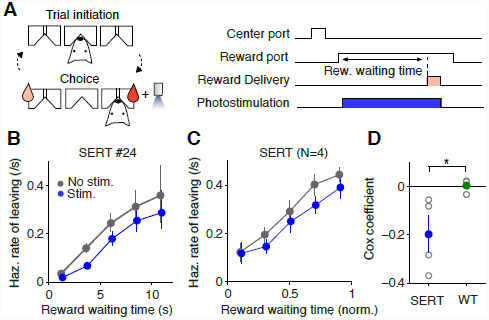
Effect of DRN 5-HT stimulation on waiting for rewards in the modified probabilistic reward task. **(A)** Schematic diagram of trial events (left) in the modified probabilistic reward task. In each trial, a mouse is required to enter the center port then move to the reward port, where it has to wait for a variable delay (min 0 s, mean selected to ensure around 50% waiting success) in order to obtain a water reward. Water drops indicate water reward probabilities associated with each side port, as in **Figure 6A**. Timeline of an example rewarded trial (right). Photostimulation was delivered from reward port entry to reward delivery (rewarded trials) or port exit (non-rewarded trials). **(B)** Hazard rate of leaving the reward port is plotted across time for an example mouse, SERT #24. Note that data from both of the reward ports (left and right) were combined. **(C)** Hazard rate of leaving the reward port for the population of SERT-Cre mice (N = 4). Averages across mice are shown. Error bars indicate S.E.M.. **(D)** Cox regression coefficients for photostimulation term are shown for SERT-Cre mice (blue) and wild-type mice (green). Individual mice shown in open circles. Averages across mice shown in filled circles. Error bars indicate S.E.M.. *: *P* < 0.05 with independent two-sample t-test (SERT v.s. WT).

## DISCUSSION

Using optogenetic tools, we selectively activated DRN 5-HT neurons while mice performed a waiting task. Our main finding was that activation of DRN 5-HT neurons resulted in an increase in mice’s ability to wait in a delayed response task but did not promote conditioned place preference, real-time place preference or choice bias in a value-based two alternative choice task.

### Serotonin promotes waiting for delayed rewards

Our waiting results are consistent with previous studies showing that reduced levels of 5-HT lead to impulsive responding for rewards [11–14, 50] (for a review, see [16]), that 5-HT neurons are active in a situation that involve waiting for delayed rewards or a delayed conditioned cue [17, 20] and that optogenetic DRN 5-HT activation prolongs waiting [19]. The present study extends these findings by showing that selective activation of DRN 5-HT neurons is not only causally sufficient to facilitate ‘patient’ waiting, but that this modulation is independent of appetitive or aversive affects, assessed via changes in preference and choice.

As with similar gain-of-function experiments, it is possible that DRN 5-HT photostimulation produced non-physiological firing rates or that photostimulation could paradoxically inhibit DRN 5-HT neurons due to depolarization block [51]. We believe these possibilities are unlikely. First, previous electrophysiological studies reported tonic activation of (putative 5-HT) DRN neurons during waiting in a similar task [17]. Thus, we are modulating 5-HT neuron activity while these neurons are likely already active. Second, previous slice experiments demonstrated that 5-HT neurons monotonically increase their firing rate up to 20 Hz photostimulation frequency [38]. Finally, we observed a monotonically increasing dose-dependent change in waiting performance using 1 to 25 Hz – a range that includes frequencies observed in previous DRN electrophysiological studies [17, 22, 23]. Therefore, it is likely that the behavioral effects we report here reflect enhancement of physiological function of 5-HT neurons.

It is also possible, as with similar optogenetics experiments, that our effects are enhanced by a relatively high degree of synchronization produced by photostimulation. However, Miyazaki *et al*. [19] showed that prolongation of waiting can be produced using a step function opsin, which is unlikely to cause synchronization.

The effect of photostimulation reported here had a rapid onset, detectable within 0.5 seconds of stimulation onset using our conservative measure, and persisted throughout the stimulation period. Thus, DRN 5-HT stimulation can affect behavioral output in a sub-second time scale. This result contradicts the classical view of a slow action of neuromodulatory systems, but is consistent with relatively fast DRN neuronal responses to external stimuli [22–25].

How does 5-HT neuronal activation lead to increases in waiting behavior? One possibility is that 5-HT has a general motoric effect – that is, 5-HT activation reduces motor output in a non-specific manner and thus causes a reduction in giving up behavior. Contrary to this possibility, we found that DRN 5-HT photostimulation did not significantly affect response time to the tone or, when delivered during reward consumption, reward port staying times. It is noteworthy that the reward-port staying time was unaffected despite the apparent similarities, in terms of motor outputs, to “waiting” at the waiting port. These results indicate that 5-HT does not have a general motoric effect, inhibiting some forms of behavior (e.g. waiting) but not others (e.g. reward port staying).

Our results reinforce the notion that 5-HT’s effects depend on the type of behavioral inhibition required. In particular, 5HT has been suggested to be involved in ‘action restraint’ (ability to withhold responding until a response is appropriate), but not ‘action cancellation’ (inhibition of an ongoing response) [15, 50]. Thus, 5-HT depletion impacts the 5-choice serial reaction time (5-CSRT) task (e.g. [11, 14]), differential-reinforcement-of-low-rate (DRL) schedule tasks [52] and go/no-go tasks (e.g. [12, 53]). 5-HT depletion also increases impulsive choice (i.e., the choice of small, immediate rewards over larger, delayed rewards) in delay-discounting paradigms (e.g. [54, 55] – but see e.g. [14]) and promotes perseverative responding in reversal tasks [56, 57]. Yet, serotonin manipulations do not affect the stop-signal reaction time (SSRT) task [15, 50, 58, 59], which requires subjects to terminate an already-initiated response.

The lack of response time effect we observed is consistent with Miyazaki *et al.* [19], although we observe a tendency for mice to slow down response times when stimulated with the strongest parameter (25 Hz at 5mW). An interesting possibility is that different, or more, groups of 5-HT neurons responsible for sensory gating [38] are recruited with the strong stimulation, thus dampening the auditory processing. In contrast to the results of Miyazaki *et al*. [19], we did observe a robust effect on the movement time immediately following waiting. It is possible that our larger data set was able to resolve effects that were not evident in Miyazaki’s data or that the differences movement duration (around 0.5 s vs. around 2.5 s) and degree of training (around 2 months vs. 2 weeks) make them differently susceptible to 5-HT modulation.

Future studies will be important to more precisely determine which types of behavior are inhibited by 5-HT, and which are not, and whether the difference can be mapped on to classically defined axis such as preparatory-consummatory behaviors (e.g. [60]) or habitual/goal-directed behaviors (e.g. [61]).

### Affective processing and reinforcement

Using similar optogenetic techniques to ours, it was reported that activation of DRN 5-HT neurons itself can serve as a positive reinforcer, primarily acting via a glutamatergic input to the dopaminergic system [43]. Thus, the association of waiting behavior and 5-HT photostimulation [19] could potentially be explained as a consequence of a primary affective role. For example, if 5-HT stimulation was hedonically pleasant, increased waiting could result from mice attempting to prolong the effects of 5-HT activation.

However, four different experiments indicate that DRN 5-HT stimulation does not exert its effects on waiting through an appetitive or aversive effect. The first evidence is from Miyazaki *et al.* [19], who showed that DRN 5-HT photostimulation did not increase spontaneous nose-poking and that water amount, but not DRN 5-HT photostimulation, increased waiting time [19]. We strengthen this evidence by adding three experiments using the same stimulation parameters that produced strong and reliable waiting effects. The first two of these replicated two of the assays used by Liu *et al.* to induce reward effects. First, DRN 5-HT photostimulation did not induce conditioned place preference. CPP is a standard procedure to evaluate the appetitive properties of natural and artificial rewards and has been shown to be sensitive to pharmacological and optogenetic manipulations of the dopaminergic system [44, 62, 63]. Second, DRN 5-HT photostimulation did not induce “real-time” place preference. Compared to CPP, the real-time place preference test is more similar to traditional intracranial self-stimulation experiments [64] or drug self-administration in that mice can increase the amount of exposure to stimulation. Finally, DRN 5-HT stimulation failed to bias mice’s choices in a probabilistic reward task. This test, while laborious to carry out, is advantageous as a sensitive test of possible appetitive and aversive effects: because it is probabilistic and can be run for hundreds to thousands of trials, it is adequate to study potentially small biasing effects of DRN 5-HT photostimulation.

The apparent inconsistency between ours and Miyazaki *et al.*’s results and the results of Liu *et al.*, might be explained by differences in light intensity used to stimulate DRN 5-HT neurons. While we use a maximum of 5mW, with robust waiting effects at 5mW and even detectable at 0.2 mW, Liu *et al.* used 20 mW. Other possible differences include the subtype of AAV used (AVV2/1 vs. AAV2/9), the Cre-line used (SERT-Cre vs. PET1-Cre), or differences in the precise targeting of stereotaxic injections, any of which could have resulted in a difference in the subpopulation of DRN 5-HT neurons affected. The differences are unlikely to result from differences in the temporal pattern of stimulation (phasic vs. tonic) [19, 45], since we used a similar pulsatile stimulation protocol method as Liu *et al.* [43]. Regardless of the cause, our study demonstrates that the response inhibition effects of DRN 5-HT release can be produced independently of reinforcing effects.

The fact that DRN 5-HT photostimulation does not serve as a reinforcer does not preclude what could be described as an affective role of DRN 5-HT in tasks involving patience. 5-HT neuron firing, like DA firing, appears to be evoked by cues carrying a high expected reward value [22, 24, 25, 43]. But in opposition to DA release, which invigorates actions, 5-HT would act by selectively decreasing the vigor of immediate (impulse or reflexive) appetitive behaviors, such as the approach to the reward port. It would thus promote behaviors requiring persistence, stability or inaction, as for staying at the waiting port in our task. Within the VTA DA system, different populations of DA neurons are activated by appetitive and aversive stimuli differently [65] and some DRN 5-HT neurons can be activated by aversive stimuli [33, 34]. Thus, it will be important to determine whether 5-HT can also act to potentiate waiting or inaction required to suppress immediate aversive responses [66].

### Decision circuits for waiting

How do DRN 5-HT neurons interact with other brain areas to promote patient waiting for delayed rewards? One possibility is that 5-HT is recruited to specifically inhibit behaviors that are impulsive. In this scenario, the 5-HT system would be activated whenever suppression of behavior is required. An alternative possibility is that the DRN receives and signals information about the availability of delayed benefits to downstream areas that are interpreted depending on the context. Thus, 5-HT could help animals to persistently engage in either passive or active behaviors depending on what was associated with longer-term positive outcomes. A task requiring active movement, e.g. lever pressing, to obtaining delayed rewards would help to differentiate these two possibilities.

Good candidates for downstream targets important to exert the 5-HT function of promoting waiting are brain regions involved in decision making and motor functions [67], such as medial prefrontal cortex [68], motor cortex [46] and striatum [69, 70]. A recent electrophysiology study using a similar waiting task demonstrated waiting time predictive signals in the motor cortex [46]. Photostimulation of 5-HT neurons together with monitoring activity in the motor cortex might reveal the circuit mechanism of how the activation of 5-HT neurons affects waiting behavior.

## EXPERIMENTAL PROCEDURES

### Animal subjects

Nineteen male adult C57BL/6 mice (11 SERT-Cre mice [71] and 8 wild-type littermates) were used in this study. All procedures were carried out in accordance with the European Union Directive 86/609/EEC and approved by Direcção-Geral de Veterinária of Portugal. Animals (20 – 25 g) were group-housed prior to surgery and individually housed post-surgery and kept under a normal 12 hour light/dark cycle (tested at light phase). Mice had free access to food. Water availability was restricted to the behavioral sessions, except for place preference tests in **Figure 5**. In place preference tests, mice had limited access to water for around 5 minutes outside the behavioral session. Extra water was provided if needed to ensure that mice maintain no less than 80% of their original weight. Mice performed 1 session per day, 6 or 7 days a week, except for two days of conditioning in conditioned place preference test. Experimenters were blind to the genotype of each animal throughout training, surgery and testing procedures. Eleven mice (7 SERT-Cre, 4 WT) were used in Experiment 1. A subset of these (4 SERT-Cre, 2 WT) were used in experiment 2. A different group of mice (4 SERT-Cre, 4 WT) were used in experiment 3, 4, 5 and 6.

### Stereotaxic adeno-associated virus injection and cannula implantation

Animals were anesthetized with isoflurane (4% induction and 0.5 – 1% for maintenance) and placed in a stereotaxic frame (David Kopf Instruments, Tujunga, CA). Lidocaine (2%) was injected subcutaneously before incising the scalp. The skull was covered with a layer of Super Bond C&B (Morita, Kyoto, Japan) to help stabilization of an implant. A craniotomy was performed over lobule 4/5 of the cerebellum. A pipette (20 µm inner diameter at the tip, beveled at around 45 degrees) was attached to a 5 µl Hamilton syringe (Hamilton, Reno, NV) connected to an hydraulic pump (UMP3-1, World Precision Instruments, Sarasota, FL) and viral solution(AAV2/1.EF1a.DIO.hChR2(H134R)-EYFP.WPRE.hGH, 10^13^GC/ml, University of Pennsylvania) was front-filled. The pipette was lowered to the DRN (Coordinates from Bregma: –4.7 AP and –3.1 DV) with a 33 degrees angle toward the back of the animal. The viral solution (1.2 µl) was microinjected at a rate of 100 nl/min. An optical fiber (200 µm core diameter, 0.48 NA, 5 mm) housed inside a connectorized implant (M3, Doric lenses, Quebec, Canada), was lowered into the brain, through the same craniotomy as the viral injection, and positioned 200 µm above the injection point. The implant was cemented to the skull using dental acrylic (Pi-Ku-Plast HP 36, Bredent, Senden, Germany). The skin was stitched at the front and rear of the implant. Mice were monitored until recovery from the surgery and returned to their home cage where they were housed individually. Gentamicin (48760, Sigma-Aldrich, St. Louis, MO) was topically applied around the implant. Behavioral testing started 3 weeks after virus injection to ensure good levels of expression.

### Optogenetic stimulation

Light from a 473 nm laser (LRS-0473-PFF-00800-03, Laserglow Technologies, Toronto, Canada or DHOM-M-473-200, UltraLasers, Inc., Newmarket, Canada) was controlled by an acousto-optical modulator (AOM; MTS110-A1-VIS or MTS110-A3- VIS, AA optoelectronic, Orsay, France). The AOM controlled the laser power without any auditory noise. Light exiting the AOM was collected (KT110/M, Thorlabs, Newton, NJ) into an optical fiber patch-cord (200 µm, 0.22 NA, Doric lenses), connected to a second fiber patch-cord through a rotatory joint (FRJ 1x1, Doric lenses), then to a chronically implanted optic fiber cannula through an M3 connector (Doric lenses). Laser power was calibrated for each targeted laser power using a powermeter (PM130D, Thorlabs) before and after each behavioral session.

### Experiment 1: Waiting task with waiting period stimulation

#### Apparatus

The behavioral apparatus for the task was adapted from the design developed by Zachary F. Mainen, and Matt Recchia (Island motion corporation, Nesconset, NY), originally developed for rat behavior. The behavioral control system (Bcontrol) was developed by Carlos Brody (Princeton University) in collaboration with Calin Culianu, Tony Zador (Cold Spring Harbor Laboratory) and Z.F.M. The behavioral box (20 × 17 × 19 cm, model 003102.0001, Island motion corporation), contained 3 front walls (135 degree angle between the center and the side walls) with a nose-poke port attached to each wall. The waiting port was located in the center. One of the side ports was chosen to be the reward port (counterbalanced across animals). The other port was kept inactive and inaccessible (under a black tape). Each port was equipped with infrared emitter / sensor pairs to report the times of port entry and exit (model 007120.0002, Island motion corporation). The water valves (LHDA1233115H, The Lee Company, Westbrook, CT) were calibrated to deliver a drop of 3 µl water for rewarded trials. A tone was generated by a sound card (L22, Lynx Studio Technology, Inc., Costa Mesa, CA). It was amplified (PCA1, PYLE Audio Inc., Brooklyn, NY) and presented through speakers (Neo3 PDRW W/BC, Bohlender-Graebener, Bad Oeynhausen, Germany). Blue LEDs were placed in the box ceiling and in all the ports to deliver a masking light.

#### Waiting task

Eleven mice (7 SERT-Cre and 4 wild-type littermates) were tested with this protocol. Data from 10 – 16 sessions (1 hour and 30 min each) were collected from all mice except for one mouse which lost implant at the beginning of the 5th session. Mice initiated each trial by inserting their snout in the waiting port. If the mouse “waited” (i.e. kept its snout inside the waiting port) until a tone (frequency modulated sound, 10 kHz carrier frequency, ± 2kHz maximum deviation with 10Hz sinusoidal modulation) was played, a water reward was made available at the reward port. A subsequent visit to the reward port (within 0.7 – 2.0s reward available period, variable across subjects but fixed within a subject) triggered reward delivery with a 0.2 s delay (patient trial). If the mouse moved out from the waiting port before the tone was played, no reward was available in that trial (impatient trial). The delay from waiting port entry to tone presentation was drawn randomly from an exponential distribution for each trial, with a minimum value of 0.5s and a mean value chosen for each animal, so that the mouse succeeded in waiting for the tone (patient trial) in 50% of trials. This value was fixed at the end of the training sessions. An inter-trial interval (ITI) period started after the delivery of reward (in patient trials), or immediately after reward port entry (impatient trials). If subjects did not visit the reward port within a reward available period (or equivalent period of time for impatient trials), ITI started after this period had elapsed. During ITI, a white noise was played and re-initiation of trial was not permitted. ITI ended after 3 s elapsed and the mouse had not entered the waiting port for 2 s. In order to optimize behavior, re-entrance to the waiting port during the reward available period was discouraged with a brief noise burst.

#### Stimulation protocol

Optical stimulation (a train of 10ms pulses with variable frequencies and amplitudes) was delivered when the animal entered the waiting port and terminated when the animal exited it. Thus the duration of photostimulation in each trial was equal to the waiting time of the animal on that trial. Re-entrance to the waiting port was not accompanied by optical stimulation. For each trial, stimulation frequency was chosen from five possible values [0, 1, 5, 12.5 or 25 Hz], and stimulation amplitude from three possible values [0.2, 1 or 5 mW]. The frequency and amplitude for each trial were pseudo-randomly selected so that a block of 15 trials contained trials with all the possible parameters. In order to reduce possible effect of mouse detecting visually the stimulation, masking light (blue LEDs in each port and in the box ceiling) was flashed at 25Hz in all trials (both in photostimulated and non-stimulated trials).

#### Training procedure

Training occurred in sequential stages with the time to accomplish each of the stages varying somewhat between animals. In the first stage of training, mice learned to poke in the waiting port and move to the reward port to collect reward. An LED light in the waiting port indicated that the trial could be initiated. Once the mouse entered the waiting port, the LED light in the waiting port turned off and a tone was played immediately, indicating that the water reward was available at the reward port (the left or right side port, counterbalanced across mice). An LED light at the reward port turned on once mouse exited the waiting port and turned off when a reward was delivered after reward port entry. Once the mouse showed a consistent sequence of waiting port followed by reward port entry, we moved to the second stage of training. Here we stopped using the LED light and gradually increased the delay to the tone. The increase was first made manually (1 to 2 sessions), and then automatically set to increase or decrease 0.01 s after every successful (patient) or unsuccessful (impatient) trial. When the mean tone delay reached a value passed 1.5 s, mice progressed to the third training phase, in which the tone was presented at two possible times. We started with 1.2 s and 1.6 s, and gradually increased the difference of possible times within and across sessions. Mice progressed to the next stage after showing fast and reliable responses to tone, irrespective of the time it was presented. In the final stage of training, we introduced the exponentially distributed random delay to the tone. We started with an exponential distribution with a minimum value of 0.5 s and a mean selected based on the previous stage. We let the mean change adaptively according to the performance of the animal, again with a 0.01 s increase/decrease after every patient/impatient trial. Once the mean delay became stable (i.e. did not change substantially) for at least two sessions, we fixed the value. If animals maintained the performance within a range of 50 – 60% patient trials with a fixed mean tone delay for a minimum of two days, we considered the animal professional.

After surgery, we retrained the animals and made sure the behavior became stable before testing. During this phase, we introduced the masking light and connected the optic fiber cable (with the laser turned off). Behavioral conditions at the final stage of training were identical to those of the testing sessions, except for the absence of photostimulation. Testing sessions took place at least 3 weeks after the surgery and once behavior was stable.

### Experiment 2: Reward port stimulation in the waiting task

Six mice (4 SERT-Cre and 2 wild-type littermates, subset of the 11 mice used in experiment 1) underwent this test. Data from 2 wild-type littermates are not shown here due to small sample size. This experiment was conducted in the same apparatus and task explained in Experiment 1. We collected data from 4 sessions (except for one SERT-Cre and one wild-type mouse, for which 7 sessions were collected).

#### Stimulation protocol

There were 3 block types of 30 trials each: a “no stimulation” block, where no stimulation ever occurred, a “waiting port” block, in which stimulation occurred at the waiting port throughout the waiting period in 50% of the trials (randomly selected) and a “reward port” block, in which stimulation occurred only at the reward port (photostimulation started at the reward port entry and stayed on for 3 seconds) in 50% of the trials (randomly selected).

The three block types were randomly interleaved. A train of 10 ms, 5mW photostimulation pulses were delivered at 25 Hz. Masking light (blue LEDs in each port and in the box ceiling) was flashed at 25 Hz in all trials from waiting port entry till the end of ITI.

### Experiment 3: Conditioned place preference (CPP)

Eight mice (4 SERT-Cre and 4 wild-type littermates, different from the mice used in the waiting task) underwent the CPP test. CPP was performed in a rectangular apparatus consisting of 2 side chambers measuring 18 (length) × 19 (width) × 21 (height) each, connected with a center corridor measuring 7 (length) × 6 (width). One side had black and white stripes on the walls with a grid floor, and the other had black circles on the wall and wire mesh on the floor. Mouse location within the chamber was monitored using a computerized photo-beam system and video camera (CMOSNC, SpyCameraCCTV, Bristol, UK). The position of the mouse was tracked offline using a software written in Python with OpenCV library. The CPP test consisted of 3 phases of behavioral testing over 4 days. On day 1, individual mice were placed in the center chamber and allowed to freely explore the entire apparatus for 30 minutes (pre-test). On days 2 and 3, one of the side chambers was paired with photostimulation and the other chamber was paired with no-stimulation. Mice were confined to one of the side chambers and given either optical stimulation (A train of 10 ms and 5 mW pulses at 12.5 Hz for 3 s, every 10 s), or no stimulation, for 20 minutes. Approximately 4 hours later, they were placed in the other side chamber and given the alternative treatment. Both the side of stimulation (left or right chamber) and session order (stimulation or no-stimulation treatment first) were counterbalanced across mice. On the second day of conditioning, the order of the treatments was reversed. On day 4, mice were again allowed to freely explore the entire apparatus (post-test). Masking light (10 ms pulses at 12.5 Hz for 3s, every 10 s) was delivered with a strip of blue LEDs attached at the top of the walls while animal stayed in either of the side chambers.

### Experiment 4: Real-time place preference

Eight mice (the same as CPP test) underwent real-time place preference test. The real-time preference test was conducted 4 days after the post-test of CPP. Real-time place preference was conducted in the same apparatus as the CPP test. At the beginning of the experiment, we placed the mouse in the center chamber. Mice were allowed to freely explore the entire apparatus. An entry to the stimulated chamber (the same as the chamber paired with photo-stimulation in CPP test) triggered optical stimulation (5 mW and 10 ms pulses at 12.5 Hz for 3 s, every 10 s). The train of photostimulation was repeated until the mouse exited the side chamber. Masking light (10 ms pulses at 12.5 Hz for 3 s, every 10 s) was delivered while animal stayed in either of the side chambers.

### Experiment 5: Probabilistic reward task

This experiment was conducted in the same apparatus explained in experiment 1 and 2, and using the same mice as in the place preference tests (Experiments 3 and 4). In this task, all the 3 ports (now called center port, and choice, left or right, ports) were accessible and used in the task. Eight mice (the same as preference tests) underwent a probabilistic reward task. Data from 15 sessions (1 hour and 15 min for each session) were collected from all mice.

#### Task and stimulation protocol

At the beginning of a trial, an LED at the center port was illuminated. The mouse was required to insert its snout into the center port. After the center port entry, the center LED extinguished and LEDs on each choice port were illuminated, indicating that the mouse was required to choose one of the choice ports (within a 100 s choice period). Each choice port was associated with a specific reward probability (40 or 10%) and stimulation probability (100% or 0%) for the duration of a block of trials (50 to 150 trials, randomly chosen). Reward and stimulation probabilities were never the same in the two ports (i.e. if the left port had 40% water probability, the right port had 10%, and vice-versa). There were 4 possible types of block (a combination of left high/low reward probability and left high/low photostimulation probability). The probability of each block type was not the same (25% for the block with left water high probability and left stimulation, 37.5% for the left water and right stim. block, 28.1% for the right water and left stim. block and 9.4% for the right water and right stim. block). The analysis was not affected by this difference. If the chosen side of the port was assigned to a water reward, a 3 µl of water was delivered. If the chosen side was assigned to a photostimulation, a train of 10 ms, 5 mW pulses was delivered for 1s at 12.5 Hz. If a water reward was assigned to a choice port, but that side was not chosen, the water reward remained available until that side was chosen [48, 49]. After a completion of a block of 50 to 150 trials, a new reward probability and stimulation probability was randomly chosen for the next block. The same type of block could occur in succession.

#### Training

Mice were trained in a sequence of stages with the time to accomplish each stage varying somewhat between animals. In stage 1, we trained mice to enter the center port and then move to the reward port to obtain a water reward using LED light at the port, similar to the training stage 1 of the waiting task. The water probabilities were 100% for both sides. In stage 2, we set water probability of one reward port to 90% and the other side to 0%. The water probability switched after a block of trials (100 – 200 trials, randomly chosen) without any signal. Once mice showed clear tendency to bias choice to the 90% water side and switched the bias quickly after the block switch, we moved on to the 3rd stage. Here, we gradually changed the water probabilities from the initial 90/0% probability to the final values of 40/10% in steps of 10%. The block size was also gradually reduced from 150 – 250 down to 50 – 150 trials. If a mouse showed a strong bias to one of the sides in one session, the following session started with the high water probability on the side contrary to the bias. Once mice showed reliable matching behavior (as in **Fig. 6B**, **C**), we introduced the masking lights and optic fiber cable.

#### Experiment 6: Probabilistic reward task with reward waiting

As a positive control for the effect of photostimulation in the mice tested for the conditioned place preference test, real time place preference test and probabilistic reward task, we further trained the same 8 mice in a modified version of the probabilistic reward task in the same apparatus. In this task, we inserted a delay between the reward port entry and reward delivery. Thus, the mice had to wait at the reward port to obtain water rewards. The delay was randomly chosen from an exponential distribution (minimum delay 0 s and mean delay selected at the beginning of the session to achieve ~50% of waiting success in water assigned trials based on the previous session). Because mice failed to obtain water rewards even in the water assigned trials in ~50% of trials, we used 20/80% of water reward probability, instead of 10/40% used in Experiment 5. The stimulation occurred in randomly interleaved 50% of trials. The stimulation started at the reward port entry and terminated when the mouse exited the port (or when the water was delivered in case of rewarded trials). Data from 4 sessions were collected from all mice.

### Histology

Viral expression of ChR2-eYFP and optical fiber placement was confirmed by histology after the stimulation experiments. Mice were deeply anesthetized with pentobarbital (Eutasil, CEVA Sante Animale, Libourne, France) and perfused transcardially with 4% paraformaldehyde (P6148, Sigma-Aldrich). The brain was removed from the skull, stored in 4% paraformaldehyde for more than 1 week. Sagittal sections were cut with a vibratome (VT1000S, Leica Microsystems, Wetzlar, Germany), mounted on glass slides and stained with DAPI (D9542, Sigma-Aldrich). Scanning images for YFP and DAPI were acquired with an upright fluorescence microscope (Axio Imager M2, Zeiss, Oberkochen, Germany) equipped with a digital CCD camera (AxioCam MRm, Zeiss).

### Data analysis

All data analysis was performed with custom-written software using MATLAB (Mathworks, Natick, MA). ANOVAs were performed using SPSS for Mac (version 21.0, SPSS, Inc., Chicago, IL). Sphericity corrections (Huynh-Feldt) were applied in all ANOVAs [72]. For all ANOVAs, type III sum of squares were used. Error bars represent standard error of the mean (S.E.M), unless stated otherwise. Alpha was set to 0.05.

Waiting time was defined as a time from waiting port entry to waiting port exit. Response time was defined as a time from tone onset to waiting port exit. Movement time was defined as a time from waiting port exit to reward port entry. Reward poke duration was defined as a time from the first reward port entry to the first reward port exit. In order to confirm that mice are responding to the tone, we generated a response time distribution from a shuffled data (**Fig. 1D**). We shuffled tone delays across trials and generated a response time histogram from the shuffled data. We repeated this procedure 1000 times to estimate 95 percent range of response time histograms from the shuffled data.

Change in fraction of patient trials was defined for each frequency, amplitude and mouse as a difference in fraction of patient trials in stimulated trials and fraction of patient trials in non-stimulated trials divided by fraction of patient trials in non-stimulated trials. Change in median waiting time, median movement time, median response time and median reward poke duration were calculated in the same way. Change in fraction of patient trials, median waiting time, median movement time and median response time were analyzed with a 3-way ANOVA with within-subject factors of Frequency (1, 5, 12.5, 25 Hz), Amplitude (0.2, 1, 5) and between subject factor of genotype (SERT-Cre, WT). When appropriate, 3-way ANOVAs were followed by 2-way ANOVAs for each level of genotype.

In order to assess a dose-dependent effect of frequency and amplitude of photostimulation on waiting time or hazard rate of leaving the waiting port, we used a Cox proportional hazards regression (*coxphfit* in MATLAB statistical toolbox). Briefly, we estimated the coefficients for the Cox proportional hazard model described as follows:

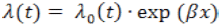

,where *λ*(*t*)represents a hazard function (hazard rate of leaving a waiting port), *λ*_0_(*t*)represents a baseline hazard function, that is a hazard function when all the covariates are 0, *β* is a row vector with 2 elements representing Cox coefficients for each covariate (frequency and amplitude), and *x* is a 2 element column vector representing covariates. Here we used frequency rank, 0, 1, 2, 3, 4 for 0, 1, 5, 12.5 25 Hz and amplitude rank, 0, 1, 2 for 0.2, 1, 5 mW as covariates. We used ranks instead of actual values based on the observation of the data (**Fig. 3A**, **C**). A coefficient of – 0.2 for frequency, for example, indicates that an increase in frequency by 1 rank (e.g.from 1 to 5 Hz), would lead to a decrease in hazard rate of leaving by a multiplicative factor of exp(–0.2) = 0.82.

Cox regression was also used to analyze reward port waiting in **Figure 4** and **Figure 7**. In these cases, we used one covariate for stimulation (0 for non-stimulated trials, 1 for stimulated trials).

Hazard rate of leaving was estimated for each time window as the number of impatient trials in which the leaving time falls on that particular window divided by the total time (summed across trials) that the mouse spent inside the waiting port during that time window. We used 5 equally-spaced, non-overlapping time windows starting at 0 and end at 95 percentile of waiting time distribution from all trials. To average hazard rate across mice, X-axis of each mice was normalized with 95 percentile waiting time. The plots for hazard rate of leaving the reward port in **Figure 7A**, **B** were plotted in the same way. For the plot of hazard rate of leaving the reward port in **Figure 4C**, we used 5 equally-spaced, non-overlapping time windows from 0 to 95 percentile of reward poke duration distribution from trials in which reward poke duration was shorter than 3 s of stimulation periods.

Nelson-Aalen estimator of cumulative hazard rate function,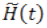, was calculated as follows(http://en.wikipedia.org/wiki/Nelson-Aalen_estimator):

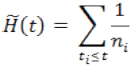

, where *t*_*i*_ denotes the time of an impatient leaving event and *n*_*i*_ denotes number of trials with waiting time more than or equal to *t*_*i*_.

To calculate the onset of the photostimulation effect, we used trials with the best photostimulation parameters (the lowest Cox regression coefficient when Cox regression analysis was performed for non-stimulated trials and trials stimulated at one particular amplitude/frequency combination, using only one covariate for with (1) or without (0) stimulation). First, we obtained the delta cumulative hazard (10 ms resolution) by subtracting cumulative hazard in non-stimulated trials from cumulative hazard of stimulated trials. Next, we defined time points with delta cumulative hazard significantly below zero as those time points in which the delta cumulative hazard was below 5 percentile of delta cumulative hazard of the shuffled data (shuffling stimulated and non-stimulated trials, 1000 iterations). We defined detectable onset time as the first time point of a number of successive points (above a criterion number) in which the delta cumulative hazard was significantly below zero. The criterion number was defined as 95 percentile of the maximum number of consecutive time bins where delta hazard of shuffled data was significantly below zero. This is a conservative measure of the onset and the true onset time is likely to be earlier than the time detected with this method.

In order to assess preference for two side chambers, we calculated preference score defined as difference between time spent in stimulated chamber and time spent in non-stimulated chamber divided by the sum of two. To test whether the mice were exhibiting proper matching behavior [47], we calculated income ratio as the amount of water obtained at the left port divided by total amount of obtained water in the past 20 trials, and choice ratio as the fraction of left choices in trials grouped (in 10 equally sized bins) according to the income ratio of the past 20 trials. In order to test an effect of the probability of water reward on choice behavior while keeping the stimulation effect constant, we calculated the probability of choosing the stimulated side [P(stim. side choice)] separately for blocks in which the higher water probability side was at the same side as the stimulation side [P(stim. side choice) for “same”] and for blocks in which the higher water probability side was opposite to the stimulation side [P(stim. side choice) for “opposite”]. P(stim. side choice) for “same” was an average of P(stim. side choice) from the two block types in which the higher water probability and stimulation were at the same side (both at the left or both at the right port). Similarly, P(stim. side choice) for “opposite” was an average of P(stim. side choice) from the two block types in which the higher water probability and stimulation were at opposite sides.

In order to test an effect of photostimulation on choice behavior while keeping the water reward effect constant, we calculated the probability of choosing the higher water reward side [P(higher water side choice)] separately for “same” blocks and “opposite” blocks.

We also analyzed the mice’s choice behavior using a logistic regression analysis [49]. A logistic regression model was estimated to predict choice on the current trial using the past trial history. The model can be formalized as:

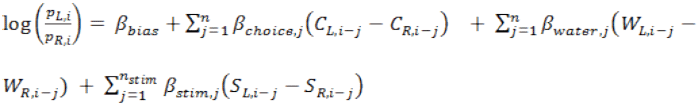

, where *p*_*L,i*_ is probability of choosing left port, *C*_*L,i*_ for a left choice (1 for choice left,0 for choice right), *W*_*L,i*_ for water obtained at the left port (1 for obtaining water at left, 0 otherwise) and *S*_*L,i*_ for photostimulation at the left port (1 for photostimulation received at left, 0 otherwise) on *i*th trial. Subscript “R” indicates the right side. βcoefficients are the parameters to be estimated.*β*_*bias*_ is a coefficient for a bias term, *β*_*choice,j*_ is a coefficient for choice, *β*_*water,j*_ is a coefficient for obtained water, and *β*_*stim,j*_ is a coefficient for photostimulation on the *j*th trial back. Maximum likelihood estimate was used to estimate *β* coefficients. We took into account up to *n* = 20 trial back. Because between nearby trials are highly correlated due to 100% or 0% of photostimulation probability in each block, we only used up to *n*_*stim*_= 1 trial back for the model presented in the main figure. Thus, total 42 parameters were estimated. Including up to *n*_*stim*_= 20 did not change the result (none of the mice showed *β*_*stim,1*_ significantly different from 0, data not shown). In the main figure we only plotted up to the 10th trial back for visualization purpose.

## AUTHOR CONTRIBUTIONS

M.S.F., M.M. and Z.F.M. designed the experiments, analyses and wrote the manuscript. M.S.F. conducted the experiments with assistance from M.M.. M.S.F. and M.M. analyzed the data.

## ACKNOWLEDGEMENTS

We thank the Mainen laboratory, in particular Eran Lottem, for many discussions; Leo Madruga, Ana Santos, Susana Dias and Enrica Audero for technical assistance; Joe Paton, Rui Azevedo and Marina Fridman for help setting up the probabilistic reward task; and Ricardo Ribeiro for help with mouse tracking. We also thank Bassam Atallah, Eric DeWitt, Eran Lottem, Dhruba Banerjee and Joe Paton for helpful comments on the manuscript. This work was supported by Fundação para a Ciência e a Tecnologia (SFRH/BD/52446/2013, M.S.F., SFRH/BPD/46314/2008, M.M.), European Research Council Advanced Investigator Grant (250334, Z.F.M.) and Champalimaud Foundation (Z.F.M.).

